# A Biphasic Approach for Characterizing Tensile, Compressive, and Hydraulic Properties of the Sclera

**DOI:** 10.1101/2020.05.21.108936

**Authors:** Dillon M. Brown, Machelle T. Pardue, C. Ross Ethier

## Abstract

Measuring the biomechanical properties of the mouse sclera is of great interest, since altered scleral properties are features of many common ocular pathologies, and the mouse is a powerful species for studying genetic factors in disease. Here, a poroelastic material model is used to analyze data from unconfined compression testing of both pig and mouse sclera, and the tensile modulus, compressive modulus, and permeability of the sclera are obtained at three levels of compressive strain. Values for all three properties measured simultaneously by unconfined compression of pig sclera were comparable to previously reported values measured by tests specific for each property, i.e., compression tests, biaxial tensile tests, and falling-head permeability assays. The repeatability of the approach was evaluated using test-retest experimental paradigm on pig sclera. Repeatability was low for measured compressive stiffness, indicating permanent changes to the samples occurring after the first test. However, reasonable repeatability for tensile stiffness and permeability was observed. The intrinsic material properties of the mouse sclera were measured for the first time. Tensile stiffness and permeability of the sclera in both species were seen to be dependent on the state of compressive strain. We conclude that unconfined compression testing of sclera, when analyzed with poroelastic theory, can be used as a powerful tool to phenotype mouse scleral changes in future genotype-phenotype association studies.

**Statement of Significance:** Ocular biomechanics is strongly influenced by the sclera, the outermost white coat of the eye. Many ocular diseases are believed to be influenced by pathological changes to scleral microstructure and biomechanics, making intrinsic biomechanical properties an important outcome measure in many studies. However, the small mouse eye precludes the use of most traditional biomechanical characterization techniques. Here, we show that unconfined compression testing analyzed with poroelastic theory can produce measurements of biomechanical properties in the pig sclera comparable to those measured by other traditional techniques. Importantly, this technique can be successfully applied to the mouse sclera, enabling more widespread use of the species as a model for ocular disease.

## 1. Introduction

The eye captures, refracts, and senses light, and the sclera, the outer white coat of the eye, has an important role in supporting these three functions. Specifically, the sclera resists the stresses imposed on the eye wall by the intraocular pressure, making its structure and biomechanical properties critical to overall ocular shape and size [1]. Scleral biomechanics also directly impact the biomechanical loading imposed on the other more delicate tissues of the eye, including the retina and optic nerve head [2–4]. Pathological remodeling and altered material properties of the sclera have been implicated in a number of ocular pathologies, including myopia and glaucoma [5–7]. Thus, more research on scleral biomechanics and how scleral microstructure influences its biomechanics is warranted.

The sclera is made up of collagen, proteoglycans, water, ions, and cells. The mammalian sclera consists of a hydrated, highly collagenous solid matrix sparsely cellularized by scleral fibroblasts. The collagen, primarily type I, is organized into lamellae embedded in a proteoglycan matrix. A similar matrix has been found to impart complex triphasic biomechanics to articular cartilage by influencing hydration and ion distributions throughout the tissue [8], and it is likely to generally function in a similar capacity within the sclera. The proteoglycan-collagen matrix imbibes a large amount of fluid, making sclera ∼70% water by weight [2]. Scleral fibroblasts actively maintain this proteoglycan and collagen matrix; however, a subset of these cells have also been shown to have contractile abilities, allowing them to interface with and actively influence biomechanics [9].

Despite its microstructural complexity, most biomechanical studies on the sclera treat it as a uniphasic material and apply linear elastic, hyperelastic, or viscoelastic material models to characterize the tissue. These studies have primarily been concerned with determining scleral response to tensile loads through uniaxial, biaxial, and inflation tests [5,10,11] and are generally performed in species well suited to ocular biomechanical studies due to the large size of their eyes e.g. pigs, cattle, and monkeys [12]; unfortunately, these same species are poorly suited to studying connections between microstructure, biomechanics, and molecular factors due to a relative absence of molecular reagents and difficulties in directly manipulating genetic factors. On the other hand, the mouse is well suited to molecular studies; however, current methods of characterizing tissue-level material properties do not scale down well to the small mouse eye (∼3mm diameter), and thus, the mouse has only been involved in a handful of ocular biomechanics studies [13–16].

Our goal was to develop methodology for quantifying the material properties of the mouse sclera. To do so, we took advantage of the microstructural complexity of the sclera and performed unconfined compression experiments analyzed with a biphasic poroelastic model originally developed in cartilage. We show that this approach can reliably characterize material properties of porcine sclera and obtain results comparable to other methods in the literature. We further show that compression testing is able to biomechanically interrogate the much smaller mouse sclera and report its intrinsic material properties for the first time. The ability to accurately measure the properties of the mouse sclera is desirable as it allows for a more direct exploration of genetic contributions to scleral biomechanics and associated ocular disorders through the use of transgenic animals and other technologies.

## 2. Materials and Methods

### 2.1 Animals

All animals were treated according to the Association for Research in Vision and Ophthalmology (ARVO) Statement for the Use of Animals in Ophthalmic and Vision Research and relevant Institutional Animal Care and Use Committee-approved protocols at the Atlanta Veteran Affairs Healthcare System. Male C57BL/6J mice were ordered from Jackson Laboratory (Bar Harbor, Maine, USA) and housed on a 12 hr:12 hr light:dark cycle in the Atlanta Veterans Affairs Healthcare System animal facility. Food and water were provided to the mice *ad libitum*. A total of 7 mice from 3 litters were used in the study. Mice were sacrificed by CO_2_ asphyxiation and eyes were immediately enucleated, marked for orientation, and stored in 0.1M (1X) PBS. Porcine eyes were obtained from a local abattoir (Holifield Farms, Covington, GA) and stored in a solution of HBSS (MP Biomedicals) with antibiotics. All eyes were maintained at 4°C until use and used the day of animal sacrifice, except where otherwise stated.

### 2.2 Sample Preparation

Porcine eyes were dissected in open air with periodic application of PBS to prevent dehydration; mouse eyes were dissected while submerged in PBS. Orbital muscle, fat, and conjunctiva were cleaned from the outer coat and then eyes were bisected at the corneoscleral junction, and the intraocular tissues were removed to create a scleral shell. This shell was then opened to create four oriented regions and the choroid and retinal pigment epithelium were removed by gently scraping with angled forceps **(Fig. 1)**.

**Fig. 1.**
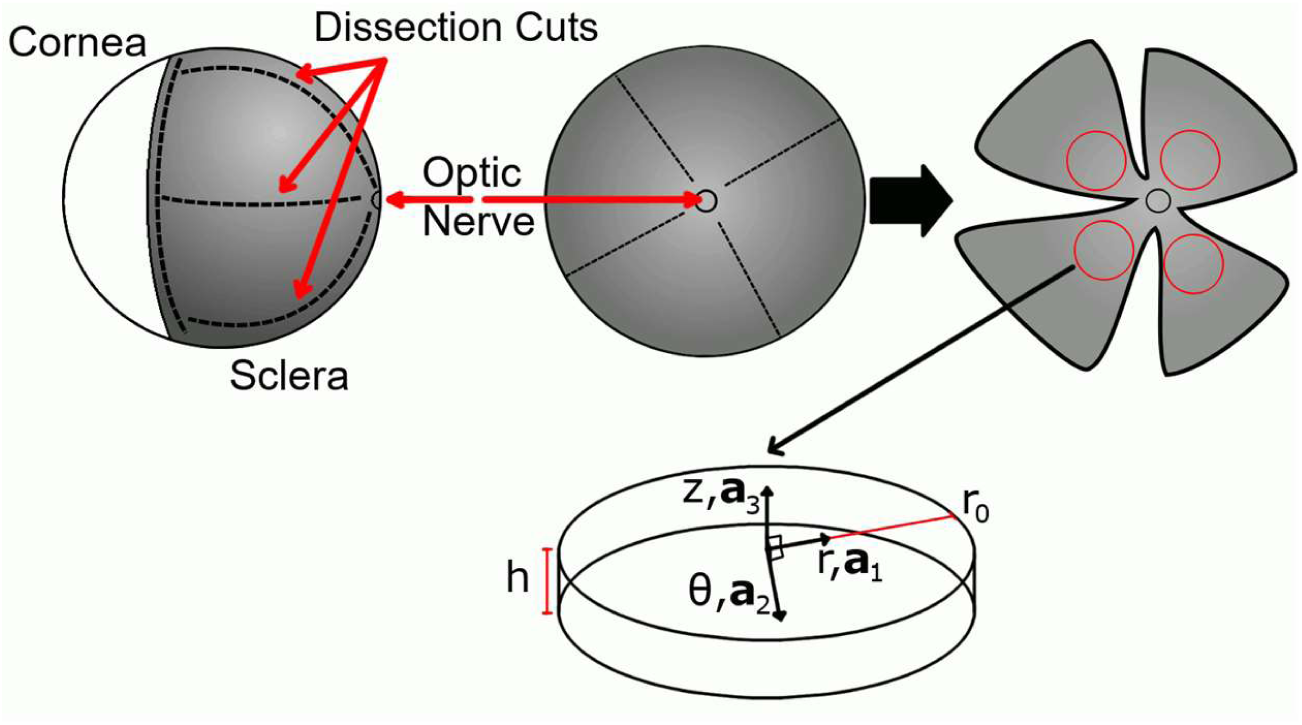
Schematic detailing the dissection procedure in a mouse eye. Top: Cornea was removed at the corneoscleral junction and intraocular tissues were removed. The resulting scleral shells were opened into four regions. Red circles show the typical punch locations in the mouse eyes; dashed lines show dissection cuts. Pig eyes were dissected similarly. Bottom: Punched samples were assumed to be cylindrical and a cylindrical coordinate system was assigned as shown. Scleral samples were always taken such that the axial cylindrical axis z and the material symmetry axis **a**_**3**_ were coincident. r_0_ and h are the sample radius and thickness in the reference configuration.

A 1mm diameter sample was harvested from the posterior region of the scleral shell using a biopsy punch immediately prior to beginning a compression test. Porcine and mouse samples were taken approximately 5mm and 0.5mm away from the optic nerve head, respectively. This location was chosen to avoid excessive circumferential alignment of collagen around the optic nerve head while remaining primarily in the posterior sclera [2]. Graphite powder was applied to scleral samples to reduce friction between samples and compression platens.

Eyes were rejected prior to testing if they were grossly abnormal, *e.g.*, had cataracts or evident damage to the sclera from the enucleation or dissection. Samples were rejected prior to testing if the biopsy did not cleanly remove the sample. Due to the scale of the mouse scleral samples, these samples additionally were rejected if folding of the sample was observed.

### 2.3 Unconfined Compression

A cantilever-based compression testing apparatus (CellScale Microsquisher, Waterloo, ON, Canada) was used to perform unconfined axial compression tests on scleral samples. This device incorporates a microcontroller that directly manipulates the position of the base of a cantilevered tungsten beam with known geometry, and a calibrated camera measures the beam tip position (resolution: 0.6155 um/pixel). A glass platen is fixed to the beam by cyanoacrylate adhesive **(Fig. 2)**. Software (Squisherjoy v5.34) calculates the deflection of the beam, which is then used to calculate the force applied to the sample in real time throughout the test. Data was collected at the compression system’s maximum sampling rate of 5Hz.

**Fig. 2.**
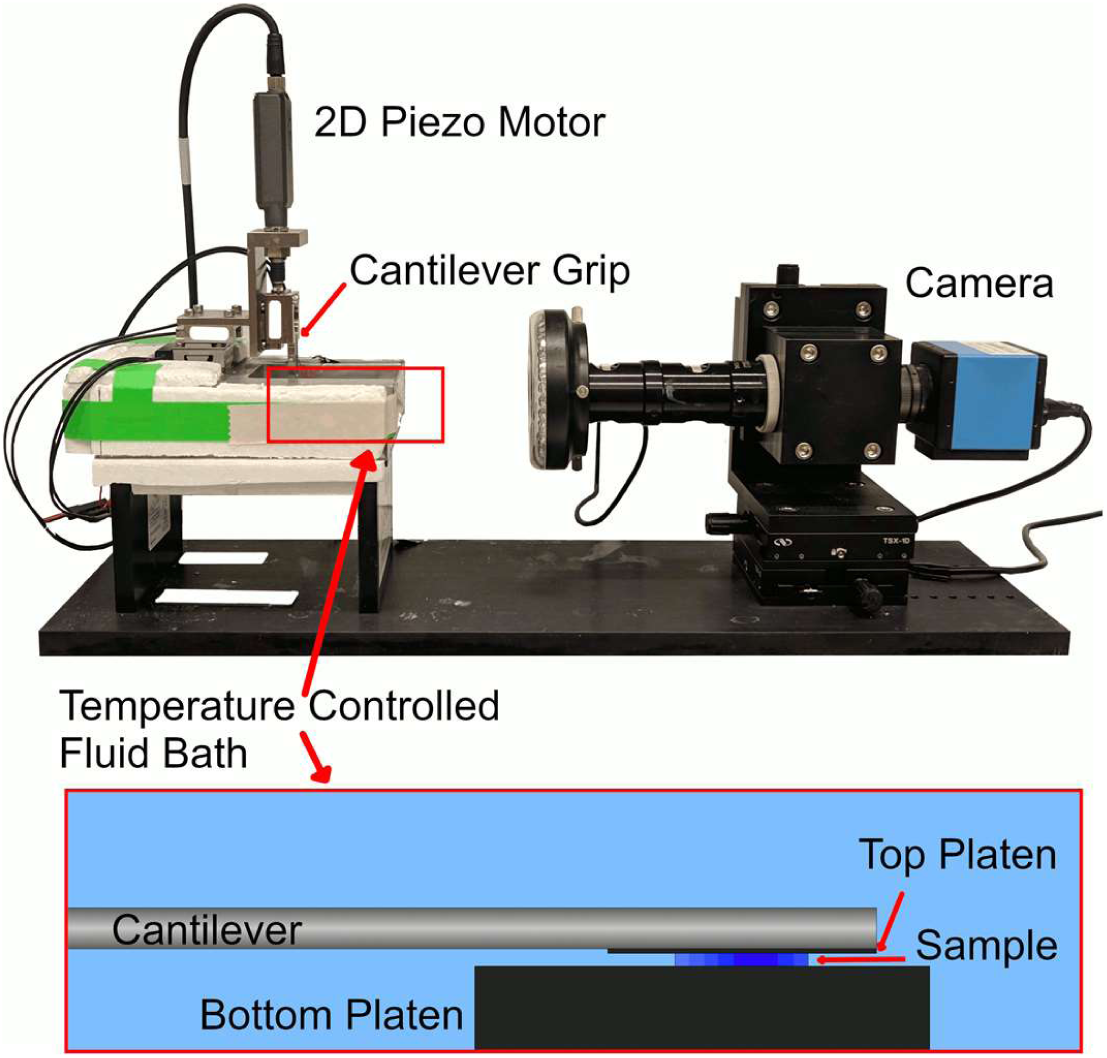
CellScale Microsquisher Compression Device. Top: Side view of the device. A piezo motor controls the gripped base position of a cantilevered beam. Tip position of the beam is measured by a camera and image processing software. Bottom: Side view of the interior of the fluid bath. The top platen is affixed to the cantilevered beam by cyanoacrylate adhesive.

Both pig and mouse samples were submerged in a temperature-controlled bath set to 37°C and underwent the same short cyclic preconditioning protocol **(Fig. 3)**. After preconditioning, a 500 μN (636 Pa engineering stress) tare load was applied to flatten the samples and ensure good contact between the sample and platens prior to stress relaxation, as is standard in compression tests [17,18]. Samples were allowed to creep under this load until equilibration (∼30 minutes). In preliminary experiments on mouse sclera where only the tare loading was performed, the time taken to equilibrate under the tare load was variable, which likely contributed additional variability in measured scleral material properties. The preconditioning protocol was empirically selected so that loads remained below physiological compressive stresses applied to the sclera by the intraocular pressure (∼2kPa), while reducing the variability in equilibration time after tare loading and in fitted parameters. Similarly, the magnitude of tare loading was deemed appropriate, since it was comparable to typical values previously used [18,19], was less than the physiological intraocular pressure, and the total applied compressive stress at final equilibrium (including the tare load) reached approximately physiological intraocular pressure (2.16±0.31kPa). The thickness of each sample at equilibrium was measured optically using the testing device’s calibrated camera. The equilibrated configuration was used as the reference for calculations of stress and strain.

**Fig. 3.**
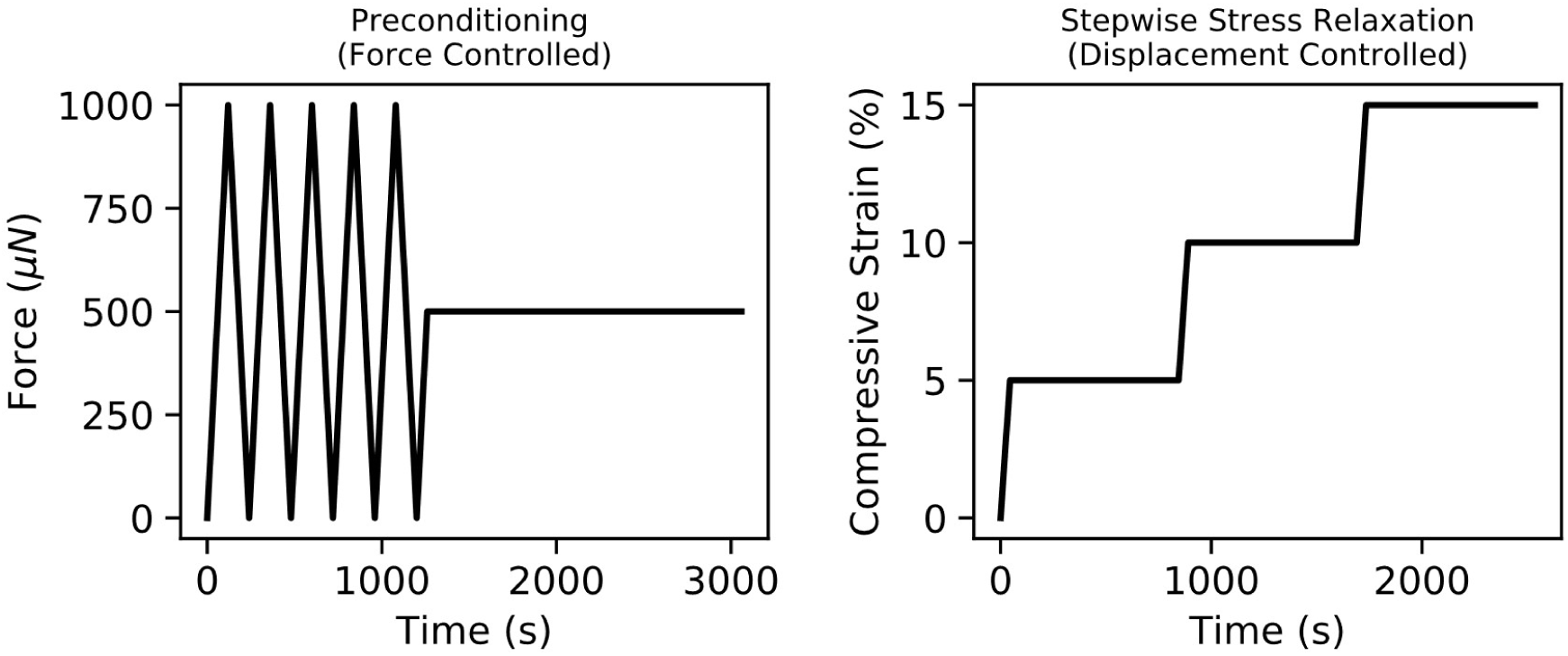
Preconditioning and stress relaxation. Left: Preconditioning protocol applied to all samples under force-controlled mode of the compression device. Right: Representative stepwise stress relaxation experiment applied under displacement controlled mode. Exact time of relaxation and ramp time varied by sample.

Immediately following equilibration of tare loading, stepwise stress relaxation experiments were performed. 15% compressive strain was applied over three steps, allowing stress to relax and equilibrate before initiating the subsequent step. 15% compressive strain was chosen as the maximum applied strain in order to conform to the assumption of small strains used to formulate the model. Practically, it was also very difficult to compress the thin mouse sclera beyond this point due to the bending beam contacting the bottom platen. Mouse and porcine sclera were compressed at ∼0.01 s^−1^ and 0.001s^−1^, respectively. The compression testing device is only capable of directly driving and controlling the base of the beam; however, sample displacement and strain is dependent on the tip displacement. Thus, the device uses software to control the tip position and perform displacement-controlled tests, which greatly limits the accuracy and consistency of applying the commanded strain. In many cases, experimental strain transiently overshot the setpoint strain significantly; however, in all cases, equilibrated strain fell within 0.06% of setpoint strain. Samples were rejected after testing if all three strain steps were not successfully applied (3 samples), if the cyanoacrylate glue failed to sufficiently fix the glass platen to the beam throughout the entirety of the test (3 samples), or if significant stress relaxation was still occurring when the next step in an experiment began (1 sample).

Porcine samples were used to determine the repeatability, or test-retest reliability, of the experimentally measured scleral material properties. After the initial stress relaxation test, porcine samples were stored in PBS at 4°C until retesting the following day. While a perfect test-retest scenario is near-impossible when working with biological tissue, since the samples may change from test 1 to 2, the concordance of the two measurements helps to inform understanding of the noise floor of the experimental method when determining material properties. Mouse samples were not used for test-retest studies due to their extremely small size, which made extended handling difficult due to their tendency to fold on themselves. Very precise alignment of the compression device was also required for mouse samples, which made obtaining two fully successful experiments on the same sample challenging.

### 2.4 Biphasic Poroelastic Theory

A modified formulation of the Biphasic Conewise Linear Elastic (BCLE) theory of Soltz and Ateshian [17] was used to model the unconfined compression response of cylindrical scleral samples. The theory considers a material composed of two incompressible phases, namely a solid matrix and an imbibing fluid. The solid matrix is postulated to be bimodular, i.e., it responds differently to tension and compression. Major points of the derivation are included below for completeness, closely following the development laid out by Soltz and Ateshian.

The total second Piola-Kirchoff stress ***S*** experienced by a biphasic poroelastic material can be decomposed into the stress that results from the solid matrix deformation, ***S***^*e*^, and an isotropic normal stress due to the interstitial fluid pressure,

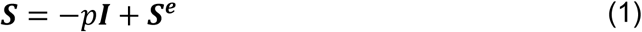

Two constitutive relations are assumed. First, fluid flux in the tissue is linearly related to the pressure gradient, i.e. Darcy’s law holds

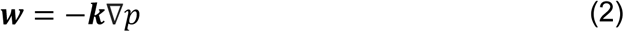

where ***w*** is the flux of the fluid relative to the solid, ***k*** is the permeability tensor, and ∇*p* is the pressure gradient.

Second, Conewise Linear Elasticity (CLE) of Curnier et al [20] is assumed for the solid matrix. The strain-energy density function, *W*(***E***), for the octantwise linear orthotropic model can be expressed as

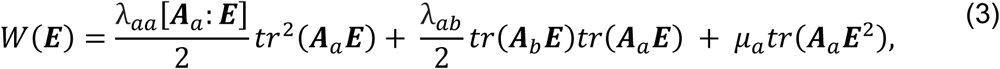

which relates the stress tensor ***S***^*e*^ to the Green-Lagrange material strain tensor ***E*** by

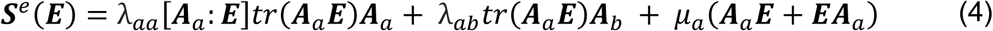

for *a, b* = 1,3; *a* ≠ *b*, where λ_*ab*_ and *μ*_*a*_ are the elastic and shear constants, ***A***_***a***_ are three texture tensors that define the unit normals describing three orthogonal hyperplanes of material symmetries, ***A***_***a***_ = ***a***_***a***_ ⨂ ***a***_***a***_, and *tr* () is the trace operator. This model captures tension-compression nonlinearity through the discontinuity of the elastic constants across tensile and compressive deformations in the three orthogonal directions ***a***_***a***_, represented by

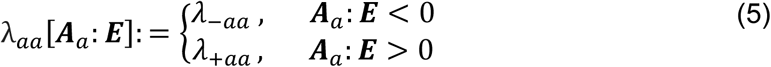

which leads to 9 elastic constants (6 diagonal constants– a compressive and tensile modulus for each ***a***_***a***_, 3 off diagonal constants) and 3 shear constants.

Unconfined axial compression on an axisymmetric cylindrical sample, assuming frictionless contact between the platens and sample, is described by the following strain tensor

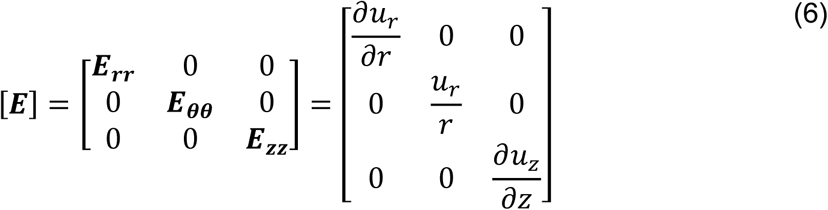

where ***u*** is the displacement vector in the current configuration, and the subscripts *r, θ*, and *z* represent the three coordinates of the cylindrical coordinate system. Under these conditions, tensile strains will be applied in the *r*- and *θ*-directions (the in-plane) and compressive strains will be applied in the *z*-direction (the through-plane).

Cubic symmetry is assumed

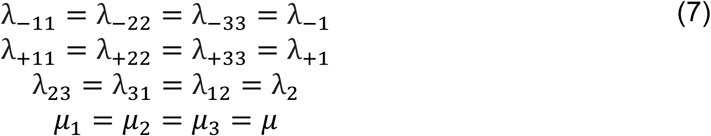

reducing the 12 material properties to 4, with the primary limitation of averaging the tensile properties into a single value. Due to the separation of tensile and compressive strains in unconfined compression of cylindrical samples, a cubic symmetry assumption is practically similar to a transverse isotropy assumption and will yield a tensile stiffness for the in-plane and a compressive stiffness for the through-plane. This assumption is reasonable for the sclera since scleral collagen fibers are preferentially aligned in-plane versus through-plane. λ_−1_and λ_+1_ can be mapped to more physically meaningful constants by the following relations

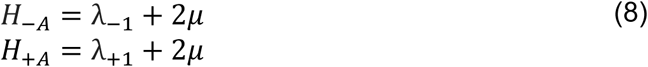

where *H*_−*A*_ is the compressive modulus of a confined sample or aggregate compressive modulus, and *H*_+*A*_ is the aggregate tensile modulus.

The stress tensor describing the matrix stresses in a stress relaxation experiment of a cylindrical sample is given by

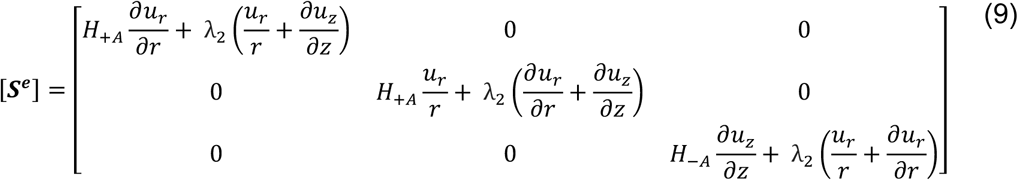

It can thus be shown that the radial component of the balance equation, neglecting body forces and inertial effects, is

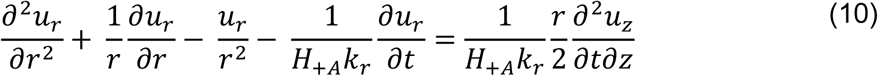

By assuming small displacements and strains and applying the appropriate boundary conditions, namely

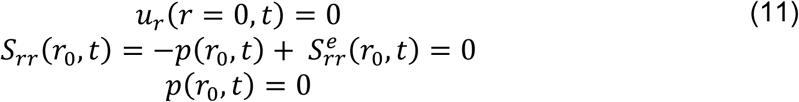

an analytical solution to the partial differential equation can be obtained in the Laplace domain, which reduces the partial differential equation to an ordinary differential equation.

Under the above assumptions of frictionless contact, axisymmetric conditions, and small strains, the axial strain imposed by the platen in a stress-relaxation experiment is independent of position and is equal to the prescribed displacement of the platens, *u*_*a*_(*t*), normalized by the thickness, *h*, of the sample,

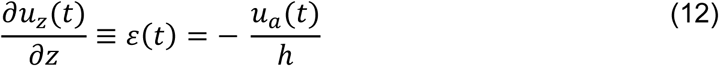

Using the radial component of Darcy’s law, the fluid pressure at any radial position can be obtained as the integral from an arbitrary radial position to the outer radius of the reference configuration, *r*_0_,

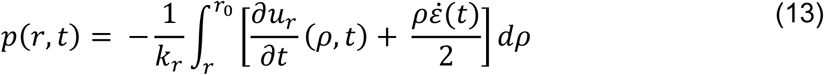

where the over dot (·) denotes a time derivative. The reaction force at the platens due to matrix deformation and fluid pressurization is thus

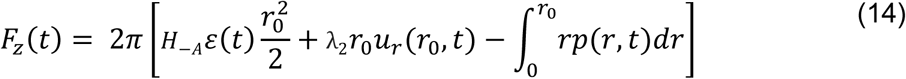

From these equations, a closed form solution can be derived to relate engineering axial stress and engineering axial strain, shown here as the dynamic modulus in the Laplace domain

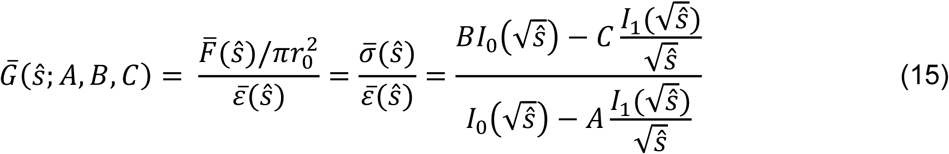

where

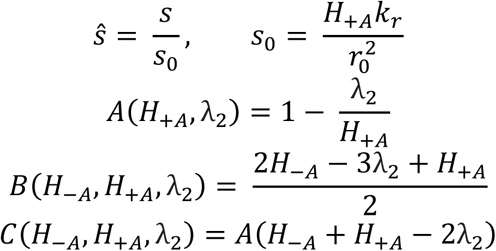

In the above expressions, *s* is the Laplace transform parameter, *s*_0_ is a characteristic angular frequency used to nondimensionalize *s, r*_0_ is the initial radius of the specimen, and *I*_0_ and *I*_l_ are modified Bessel functions of the first kind. Over bars (^−^) represent quantities in the Laplace domain and over hats (^) represent nondimensionalized quantities. The nondimensionalized dynamic modulus can further be obtained by normalizing the stress and strain signals to their equilibrated values, *σ*_*eq*_ and *ε*_*eq*_.

Four material properties fully describe the tissue under this shear-free deformation (Eq. (6)): the aggregate compressive modulus *H*_−*A*_, aggregate tensile modulus *H*_+*A*_, the off-diagonal modulus λ_2_, and the hydraulic permeability *k*_r_. The off-diagonal modulus λ_2_ can best be interpreted in the context of a confined compression experiment, where samples are physically prevented from expanding radially. If a sample were fully confined and axial compression were applied, λ_2_ would represent the ratio of stress applied to the confining walls to the amount of axial strain applied.

By taking the limit of the dynamic modulus in low frequencies (*s* → 0), we obtain a relationship between the equilibrium unconfined compressive (Young’s) modulus *E*_−*Y*_ and the other CLE constants

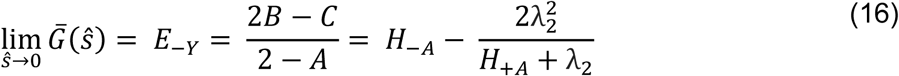

In an unconfined compression stress relaxation test, this modulus can be directly measured by taking the ratio of equilibrium stress to equilibrium strain, *E*_−*Y*_ = *σ*_*eq*_/*ε*_*eq*_. By rearranging Eq. (16) to isolate *H*_−*A*_, it is convenient to reformulate B and C to be in terms of this directly determined quantity

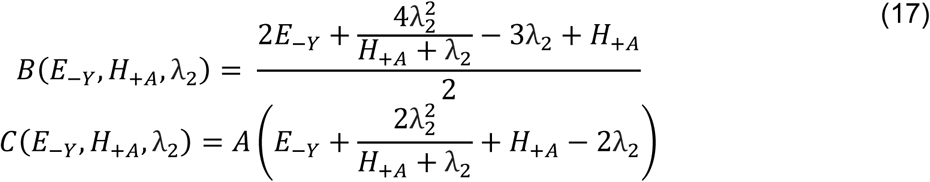

This modification to the formulation of Soltz and Ateshian allows for the direct determination of one property, reducing the dimensionality of the optimization problem when fitting the model to experimental data.

Time domain stress responses can be determined by describing the strain with an analytical function, *e.g.*, a step (*ε*(*s*) = 1/*s*) or a ramp function (*ε*(*s*) = 1/*s*^2^), and then taking the inverse Laplace transform. However, due to the large variations in applied strain in our experiments, time domain model stress was instead obtained by convolving the experimental strain data with the impulse response function *h*(*t*), given by

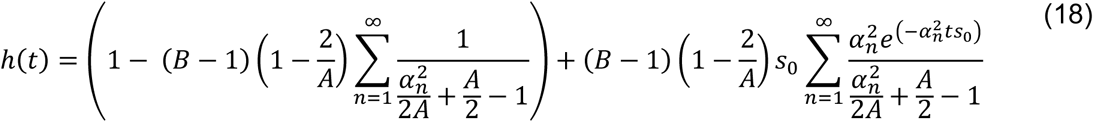

obtained by taking the inverse Laplace transform of 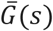. *α*_n_ are the roots of the characteristic equation describing the poles of Eq. (15)

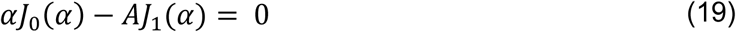

where *J*_0_ and *J*_1_ are the Bessel functions of the first kind.

### 2.5 Data Analysis

Engineering stress and engineering strain were calculated by dividing the axial force, *F*_*z*_(*t*), and axial displacement, *u*_*a*_(*t*), by the reference configuration contact area (assumed identical to the internal area of the biopsy punch) and initial sample thickness, respectively. Each of the three stress-relaxation steps for each sample were analyzed independently. In more detail, stress, strain, and time at the beginning of each step were forced to 0 by subtracting the first measurement in that step from every subsequent timepoint in the step. The final 50 seconds of each step were assumed to be fully equilibrated, and mean values of stress (*σ*_*eq*_) and strain (*ε*_*eq*_) over this time were used to calculate *E*_−*Y*_. Stress and strain signals were normalized by each of their equilibrated values.

Normalized stress and strain signals were each represented using a linear interpolation function specified with Heaviside step functions, which has a Laplace transform that can be analytically expressed in terms of *s*, as defined in Huan et al. [21]. Due to the inherent force-controlled nature of the compression device used, the ratio of transformed strain to stress was used to create an experimental transfer function, 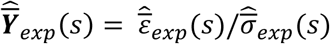, that could be used to determine the experimental gain and phase; this is in the reciprocal of the derived BCLE dynamic modulus (Eq. (15)). The final three material properties, *H*_+*A*_, λ_2_, and k_*r*_, were thus obtained by curve fitting the reciprocal of Eq. (15), 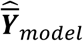, to 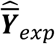 over a limited frequency range (10^−7^–10^−1^ Hz). The frequency range was informed by previous studies in cartilage and was chosen to capture the behavior of the samples while limiting the influence of measurement noise [22]. A differential evolution algorithm was used to cover a wide expanse of the parameter space and decrease the chance of local minima convergence [23] during minimization of the chosen objective function [24],

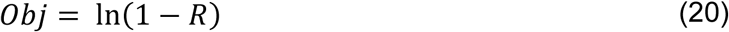

where R is the cross-correlation between the experimental transfer function and model, given by

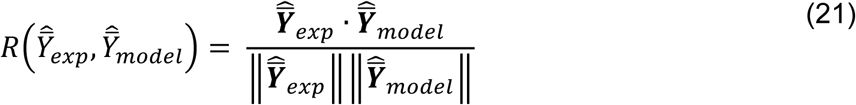

Bounds were implemented for the material properties to enforce positive definiteness of the elasticity tensor

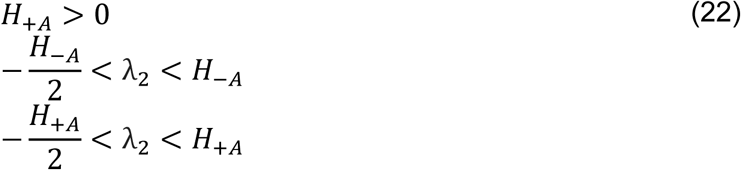

Due to the inherent dependence of permeability on compressive strain, a standard exponential model [18,25] was fit to the three permeability values obtained for each sample, given by

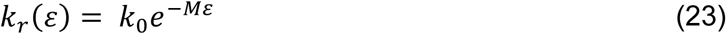

where *ε* is the compressive engineering strain, *k*_0_ represents the in-plane permeability at 0% compressive strain, and *M* defines the nonlinearity of the relationship between compressive strain and permeability.

### 3.5 Statistical Analysis

Goodness of fits of the models, 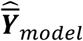, to the experimental transfer functions, 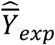, are reported as the nonlinear coefficient of determination, *r*^2^, defined as [26]

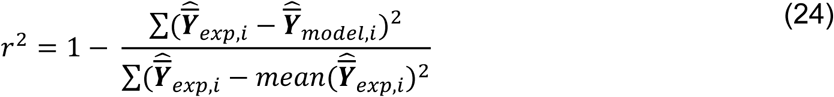

The four material properties obtained from each step of each experiment were analyzed using a random-intercept Gamma generalized linear mixed effects model (GLMM) with a log link function using the “lme4” package in R. The log link function ensures positive fitted values and is well suited to the necessarily positive tensile and compressive moduli and hydraulic conductivity. Due to the range of lambda values crossing zero, a standard shift transformation was applied to translate all values to positive for analysis. The statistical model was defined as

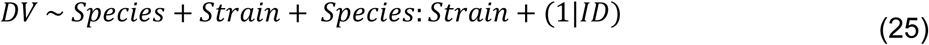

where *DV* is one of the material properties, *Species* is either pig or mouse, *Strain* is the amount of compressive strain applied, *Species*: *Strain* accounts for the interaction of the two fixed effects, and (1|*ID)* is the random intercept factor that accounts for the three repeated measurements on the same samples. This model was deemed appropriate for each material property based on the resulting normality of the residuals. The model for each material property was used to hypothesis test each effect using likelihood ratio tests between the full model (Eq. (25)) and three null models, each lacking one of the fixed effect terms. In the presence of significant interaction effects, Tukey post hoc tests were performed, and the adjusted p values are reported. Only the initial measurements of porcine sclera were included in this portion of the analysis.

The test-retest repeatability of the fitting of tensile stiffness and hydraulic conductivity and the measurements of thickness and unconfined compression modulus values between first and second test of porcine scleral samples were analyzed by calculating both Pearson (*ρ*) and Lin’s concordance (*ρ*_*c*_) correlation coefficients. Perfect concordance or discordance would be represented by *ρ*_*c*_ values of ±1, whereas a *ρ* of ±1 would indicate perfect univariate correlation.

## 3. Results

In total, 10 eyes were included for analysis (4 porcine, 6 mouse), which included 13 unique samples (4 porcine, 9 mouse) and 51 fitted stress relaxation steps (24 porcine, 27 mouse). Seven samples in total were excluded from analysis after testing. One sample was taken from each pig eye, and each sample was tested twice. When multiple samples were successfully tested from the same mouse eye, the obtained properties were averaged. All results are reported as the mean ± standard deviation unless otherwise noted. The thickness of the samples after preload was 1294±241 μm for the porcine sclera and 77.2±8.9 μm for the murine sclera.

### 4.1 Curve-fitting Quality

The BCLE theory successfully fit the unconfined compression response of scleral samples from both species **(Fig. 4)**. The average nonlinear coefficient of determination of the model to experimental data (Eq. (24)) in the frequency domain was 0.9989±0.0006 and 0.9982±0.0018 for samples from pig and mouse, respectively. Fitting was repeated ten times for each of the 51 strain steps using different starting populations selected by randomly seeded Latin hypercube sampling of the bounded parameter space. Of the 10 fits performed for each of the 51 steps, none converged on appreciably different sets of parameters. The mean standard deviation of each parameter over the repeated fittings was small (STD H_+A_: 598.0 Pa, λ_2_:150.7 Pa, k: 3.7e-16 m^4^/Ns), indicating a consistent convergence to the same set of parameters.

**Fig. 4.**
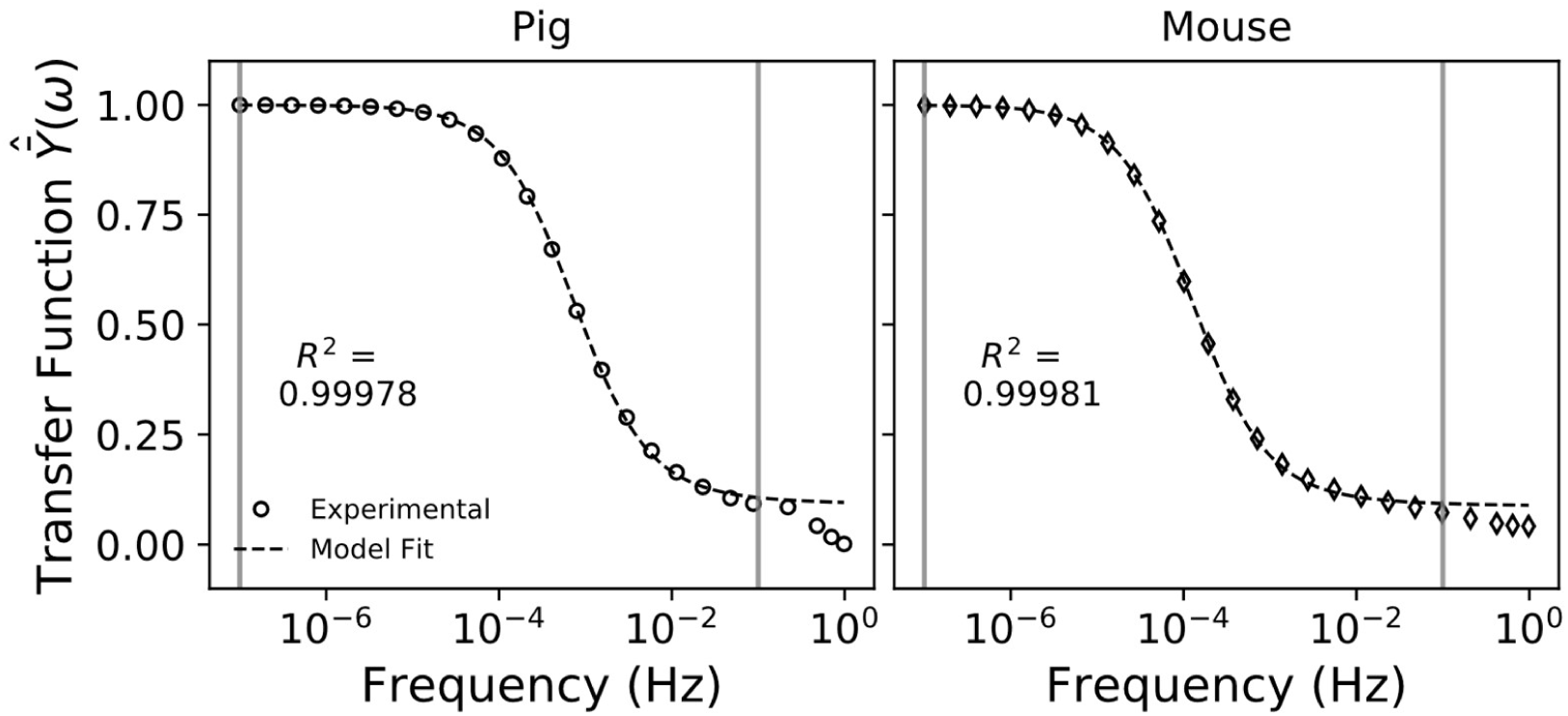
Experimental and fitted transfer functions for pig (left) and mouse (right). Vertical lines represent the bounds of the data range used for fitting. Excellent agreement was seen over fitting region as judged by the nonlinear R^2^ (Eq. (24)), with deviations present at higher frequencies. The reported R^2^ value refers only to fitting between the vertical lines.

Using the properties obtained by fitting the experimental transfer function, model phase and gain components generally agreed well with the experimental components **(Fig. 5).** The experimental gain and phase graphs for both species display the typical behaviors of biphasic materials, namely high frequency stress application results in a dampened strain response due to the fluid pressurization of the tissue, whereas slow application leads to strain magnitudes dependent on the compressive stiffness, *E*_−*Y*_. Fluid pressurization and permeation are also seen to cause a phase delay in the strain response from stress applied at intermediate frequencies, the specifics of which are dependent on the material properties. The model time domain stress obtained by convolving the experimental strain with the impulse response function also agreed well with the experimental data, showing similar peak stresses and relaxation time between model and experiment (**Fig. 6)**.

**Fig. 5.**
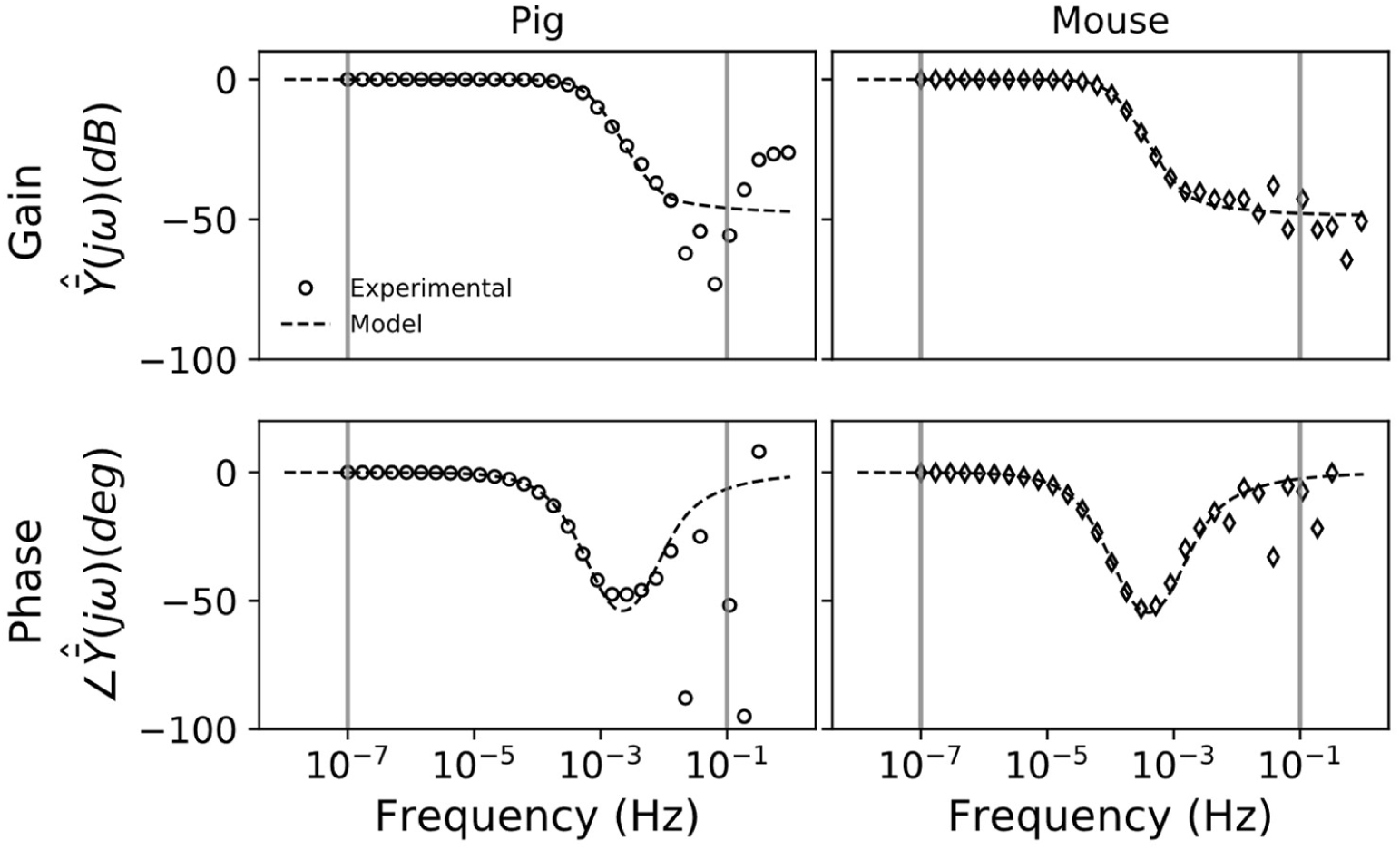
Experimental and model Bode plots for pig (left) and mouse (right) scleral samples. Top: Gain as a function of frequency. Bottom: Phase as a function of frequency. Model values were calculated using the material properties obtained by curve fitting the transfer function over a selected frequency range (represented by the vertical lines). Both species show frequency dependence for gain and phase that are matched well by the model. Noise infiltration exists at the higher frequencies primarily due to the high frequency camera noise of the device, while noise infiltrates at lower frequencies in the pig due to the slower ramp speed.

**Fig. 6.**
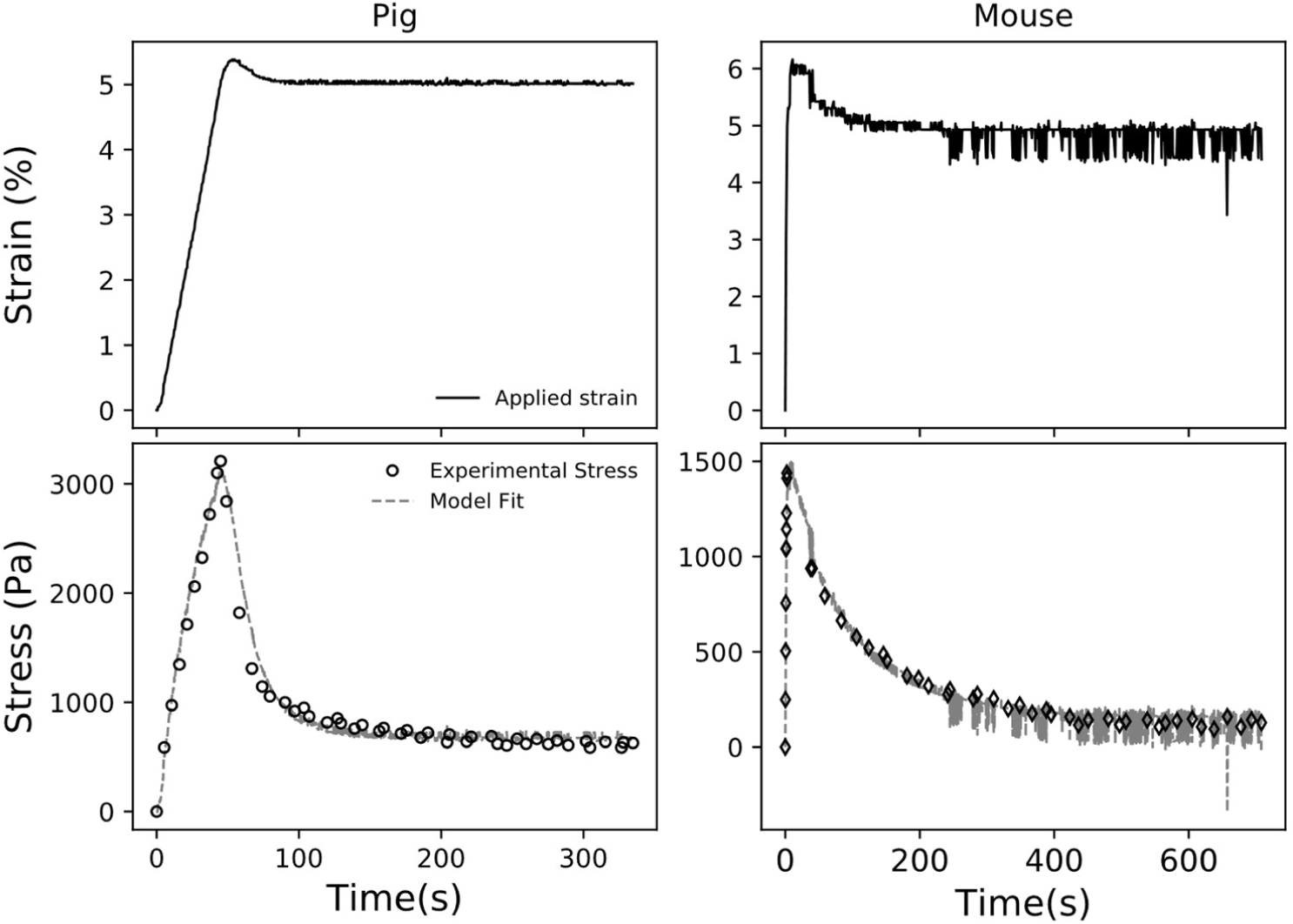
Representative strain measurements for both pig (left) and mouse (right) scleral samples at the first step (0-5% strain) and the resulting model fit. Top: Engineering compressive strain during a ramp-hold experiment at 5% compressive strain. Significant deviation from the desired strain vs. time profile is evident in both species at both ramp speeds. Bottom: Time-domain experimental compressive stress and modelled stress, as determined from the applied strain for pig and mouse scleral samples. Note, significantly more noise is visible in the mouse strain signal due to the thinner sclera. The camera and beam tracking add noise to the measurements of displacement to a similar degree in both species, but the noise makes up a smaller percentage of the overall displacements in the pig sclera.

### 4.2 Material Properties

For both species, both tensile modulus and permeability were highly dependent on the state of compressive strain of the sample **(Fig. 7)**; the tensile modulus was seen to increase with increasing compressive strain, while the hydraulic conductivity decreased, a common finding with biphasic materials. However, significant interaction effects between the state of compressive strain and species were also found for both the tensile modulus and permeability (*Species:Strain*; *H*_+*A*_: p<0.001, *k*: p<0.001), implying that both properties are dependent on state of compressive strain in a species-dependent manner. The porcine sclera was significantly stiffer in tension than the mouse sclera at the first two strain steps (p<0.001 for both 5 and 10% strain), with the difference vanishing at the third strain step (p=0.098). The murine and porcine sclera had equal hydraulic conductivities over the first two strain steps (5% strain: p=0.34; 10% strain: p=0.90), with the mouse sclera significantly less permeable than the porcine sclera at the third strain step (p<0.001). *E*_−*Y*_ and λ_2_ were not observed to change with compressive strain (*E*_−*Y*_: p=0.07; λ_2_: p= 0.29) or be species-dependent (*E*_−*Y*_: p=0.26; λ_2_ :p= 0.19) **(Table 1)**.

**Table 1.** Average material properties over strain steps and samples. n values refer to the number of eyes tested.

| Species | $E_{-Y}$ (kPa) | $H_{+A}$ (kPa) | $\lambda_2$ (kPa) | $k \times 10^{-15}$ ($m^4/Ns$) | $k_0 \times 10^{-15}$ ($m^4/Ns$) |
| --- | --- | --- | --- | --- | --- |
| Pig ($n=4$) | $10.0 \pm 3.2$ | $341.4 \pm 130.6$ | $9.5 \pm 3.6$ | $8.6 \pm 4.2$ | $25 \pm 33$ |
| Mouse ($n=6$) | $10.1 \pm 4.6$ | $173.7 \pm 118.9$ | $12.6 \pm 7.9$ | $9.5 \pm 9.7$ | $62 \pm 51$ |

**Fig. 7.**
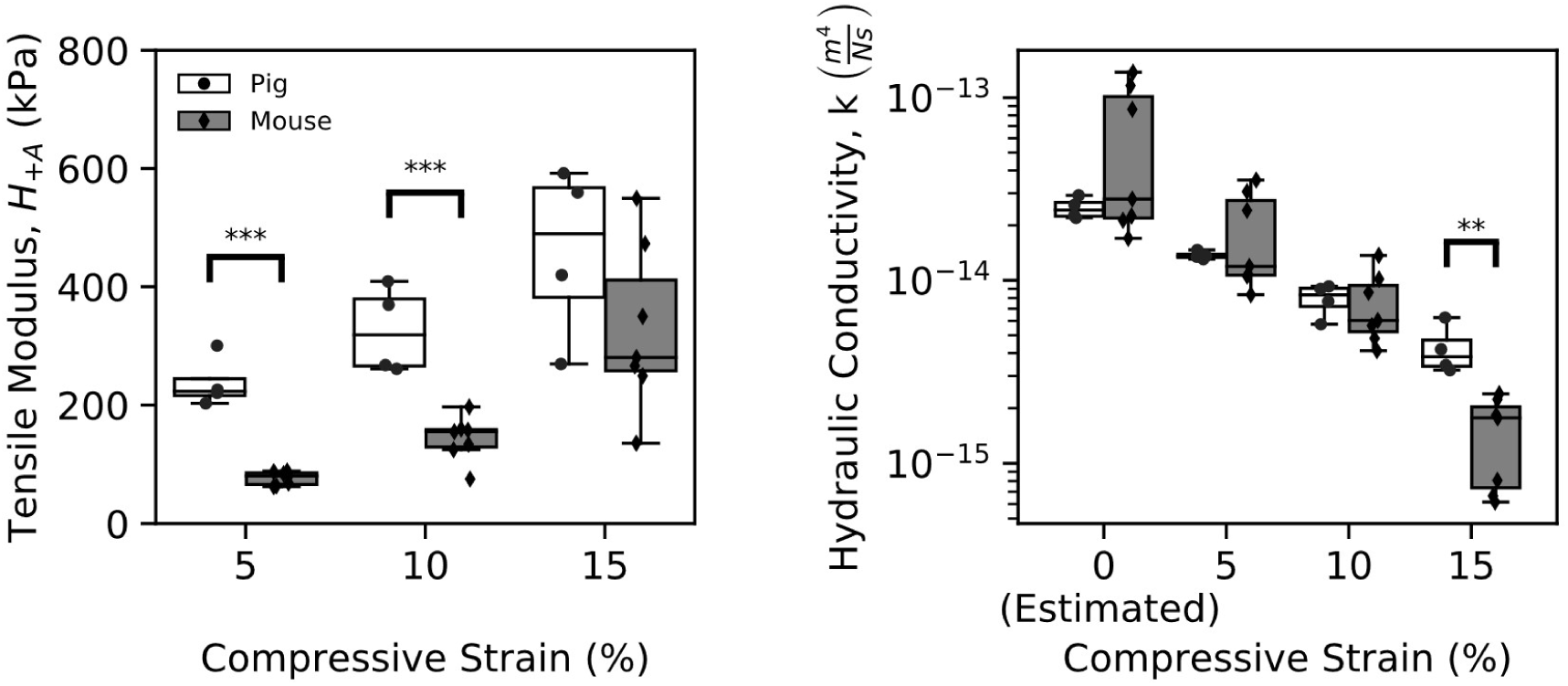
Scleral material properties as a function of compressive strain. Left: Tensile stiffness of pig and mouse sclera. Stiffness of both species’ sclerae increased with increasing compressive strain, and a significant interaction effect was present (Species:Strain; p<0.001). Right: Permeability of pig and mouse sclera. Permeability of both species decreased with increasing compressive strain, and a significant interaction was present (Species:Strain; p<0.001). ***: p<0.001, **: p<0.01 by Tukey post hoc analysis with adjusted alphas for multiple comparisons.

### 4.3 Repeatability

The measured compressive moduli and thicknesses were both highly correlated (*E*_−*Y*_: ρ=0.83, p<0.001; thickness: ρ=0.98, p=0.019); however, the concordance of the compressive moduli between experiments was only moderate (*E*_−*Y*_:: ρ_c_=0.52; thickness: ρ_c_ =0.92), showing a general trend of increasing in stiffness from the first to second test **(Fig. 8)**. The fitted tensile moduli and permeabilities were both highly correlated and moderately concordant between the first and second tests (H+A: ρ=0.82, p=0.001, ρ_c_=0.76; k: ρ=0.83, p<0.001, ρ_c_=0.82), not displaying any notable trends.

**Fig. 8.**
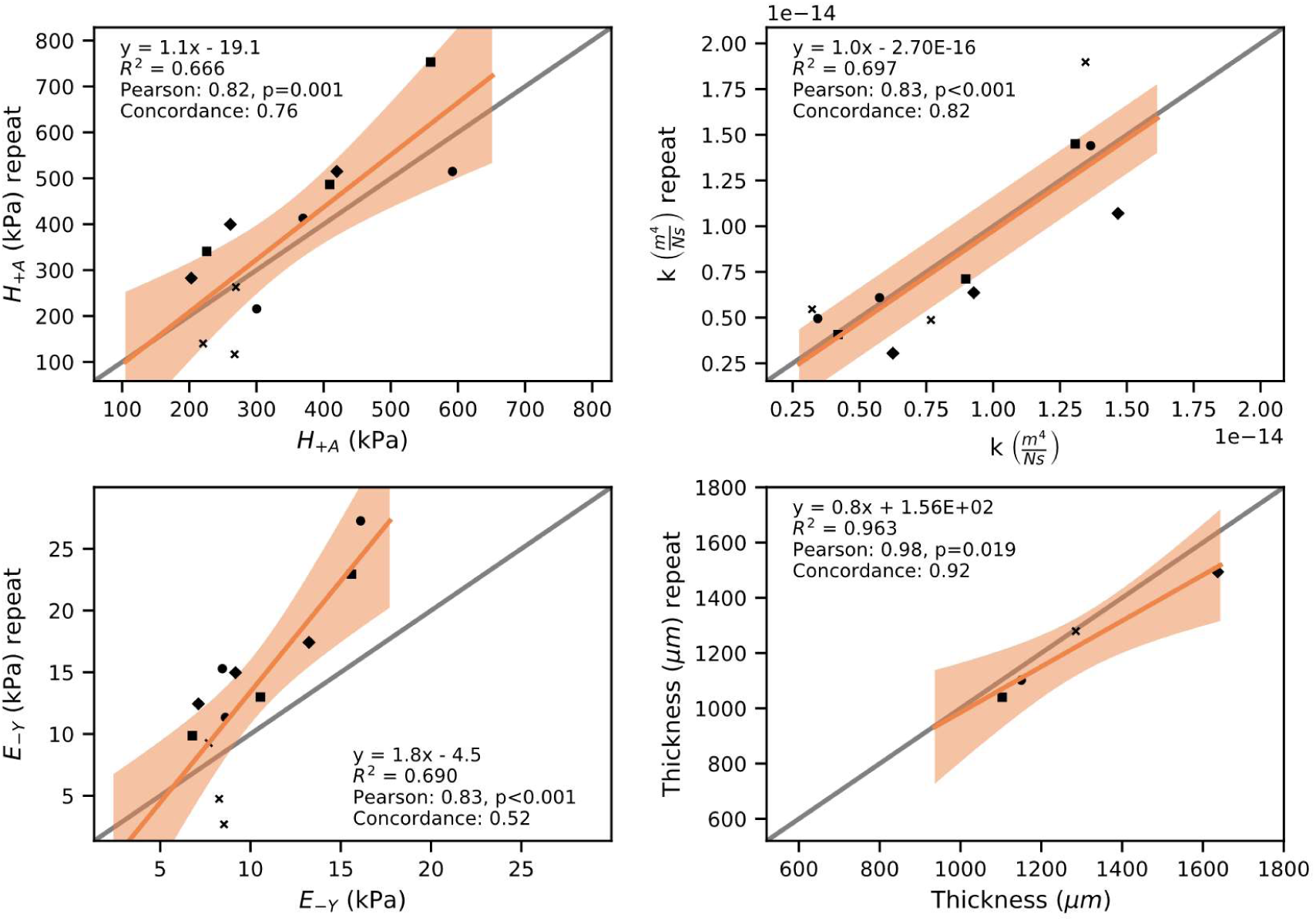
Concordance plots of first and repeated measurements on pig scleral samples. Top: Fitted properties, tensile modulus (Left) and hydraulic conductivity (Right). Both show moderate concordance and repeatability. Bottom: Directly measured properties, compressive modulus (Left) and thickness (Right). Compressive modulus is observed to change from first to second test. The identity line is represented in gray. Perfect concordance between first and second measurement would be indicated by points falling exactly on this line. Each symbol shape represents measurements coming from the same eye over the three strain steps. Orange lines show the best fit linear regression, and shaded areas show the 95% confidence bands.

## 4. Discussion

We describe here the first multiphasic biomechanical characterization of the sclera using the Biphasic Conewise Linear Elastic model and unconfined compression testing. Using unconfined compression and a closed form solution of the biphasic model, the through-plane compressive stiffness, in-plane tensile stiffness, and hydraulic conductivity of the tissue could be determined from a single stress-relaxation experiment. Importantly, good curve fitting was observed not only in porcine sclera, but also mouse sclera which was ∼15x thinner on average. This enabled us to determine the intrinsic material properties of the mouse sclera for the first time.

### 4.1 Applicability of BCLE to Characterizing Porcine Sclera

The optimal scenario for determining how well a material model captures the physical characteristics of a tissue involves fitting a measured quantity to determine tissue material properties and then using the model and fitted properties to predict another simultaneously measured quantity, *e.g.*, curve fitting stress and comparing measured and model-predicted radial expansion or interstitial fluid pressure. Due to the physical size of the samples and the constraints on the compression device, this was not achievable in this study. Thus, to gain an understanding of how well the model described the tissue response, we instead chose to compare the magnitude of the fitted porcine properties to the literature, while also judging the quality of the curve fitting and the repeatability and uniqueness of fitted properties. Each of these factors support the BCLE model as capable of describing the biomechanical response of the sclera under unconfined compression in a physiologically reasonable and unique manner, as described below.

### 4.2 Tensile and Compressive Stiffness of Porcine Sclera

The tensile stiffness of porcine sclera has been previously characterized many times using uniaxial testing; however, the in-plane strains generated from unconfined compression is biaxial in nature, and thus, comparison to uniaxial results is invalid. Biaxial testing of porcine sclera has been reported once by Cruz-Perez et al., who used a reduced Fung-type hyperelastic model to analyze biaxial tensile tests [11]. Scleral samples in their study were stretched in-plane between 2-4% and had an average equivalent stiffness of 410.81 ± 168.58 kPa. Here, BCLE theory estimates a peak in-plane tensile strain of 2.5% for any one individual step that would decrease to 0% during stress relaxation, and an overall average stiffness of 341.4 ± 130.6 kPa for porcine sclera. We conclude that despite the disparate handling, methodology, and analysis procedures between these two studies, the tensile stiffness values measured by unconfined compression matched the results of a well-accepted biaxial methodology reasonably well.

Studies on the compressive properties of the sclera are rare compared to those on its tensile properties, but the results here are comparable in magnitude to the few previous studies performed. Using unconfined compression and measuring a drained secant modulus, a conceptually similar measure of elastic stiffness to the equilibrium Young’s modulus, *E*_−*Y*_, measured in this study, Mortavazi et al. found porcine peripapillary sclera to have a drained secant compressive modulus of approximately 4kPa [27], slightly lower than the 10 kPa measured here in both porcine and mouse sclera. A study of the compressive properties of porcine ocular tissues by Worthington et al. measured a compressive Young’s modulus of sclera to be ∼35kPa, which agrees closely to what was measured in human scleral samples by Battaglioli and Kamm [19,28]. However, Battaglioli and Kamm also measured scleral samples from cattle and found the compressive modulus to be in the range of 12-18kPa, closer to the measurements found in this study. The compressive stiffness of porcine sclera measured here had a mean of 10kPa spanning ∼6.5 to 16kPa, which thus appears physiologically reasonable. The small differences in mean values between studies are likely due to the minor differences in reference configurations used for stress and strain calculations, namely differences in tare loading; however, differences in parameters used to quantify compressive stiffness (drained secant modulus, Young’s modulus, and equilibrium Young’s modulus) and regions from which scleral samples were taken could also contribute to the differences.

### 4.3 Hydraulic Conductivity of Porcine Sclera

Through-plane permeability of sclera has been measured a few times in the literature using a falling head permeability assay, but it is known that a lack of uniformity in permeability testing methodology makes comparisons between laboratories difficult [29]. Jackson et al. measured human scleral permeability and found an average value of 8.62e-18 ± 10.42e-18 m^2^ from 18 human donors [30]. Stewart et al. measured porcine scleral permeability and reported an average value of 43.3e-18 ± 23.5e-18 m^2^ from 7 pigs [31]. Due to the dependence of permeability on compressive strain and the likely anisotropic nature of the quantity due to structural anisotropy of the sclera, a direct comparison to the literature is challenging. However, after converting hydraulic conductivity measurements in this study to intrinsic permeability by assuming a viscosity of 0.7 cP for PBS at 37°C [32], the average value of porcine permeability at 5% compressive strain in this study was 9.8e-18 ± 0.50e-18 m^2^, and the estimated zero-strain permeability was 17.9e-18 ± 2.9e-18 m^2^. Thus, the measured magnitude of permeability here lies between the literature values, showing that this methodology produces a physiologically reasonable permeability value with less variability than the falling head methodology, which is prone to leakage, a lack of precisely controlled or measured compressive strain, and possible sample degradation/remodeling over the typical ∼24 hour testing times.

### 4.4 Quality of curve fitting

The model was successfully able to fit the experimental data from low frequencies up to the chosen cutoff frequency, at which point high frequency camera noise and the low sample rate of the device precluded accurate Laplace transformations **(Fig. 4)**. While the material properties were determined only from curve fitting the experimental transfer function, defined as the Laplace domain strain signal divided by the stress signal, on the real axis of the Laplace domain, both the gain (real) and phase (imaginary) components of the model calculated using these fitted properties aligned well with the experimental components. Finally, the material properties determined in the frequency domain led to time domain stress fits of similar quality; specifically, the model matched the peak stresses and characteristic relaxation times quite well after accounting for the deviation from ideal ramp-hold strain application by taking the convolution of the experimental strain with a derived impulse response function (Eq. (18)).

### 4.5 Repeatability

The ability to curve fit data alone does not prove a model’s applicability to analyzing tissue response; a model can curve fit any data given enough degrees of freedom. It is important that a model yields a unique set of parameters for any individual curve fit in order to characterize a tissue accurately and in the most generalizable manner. Here, we probed the uniqueness of the fits by utilizing a differential evolution algorithm to more fully interrogate the parameter space in conjunction with performing a test-retest experiment. The sets of parameters for any one experiment did not vary appreciably from one curve fit to another, implying that the optimization was finding a global minimum. Similarly, the sets of parameters obtained by curve fitting did not vary much from the first to second test of the samples. If the biphasic model employed here did not describe the physical state of the samples in this configuration uniquely, it is unlikely that the test-retest methodology would show concordance. A perfect test-retest experiment was not achievable, due to the likelihood of physical changes in the samples between the first and second test either due to the loading experienced in the first test, the time spent in PBS overnight, or both. Even though the directly measured compressive modulus of the samples were seemingly altered slightly from the first to second test, as shown in the lack of concordance, the stiffness and hydraulic conductivity obtained through curve fitting were both still reasonably concordant.

### 4.6 Murine vs Porcine Sclera

Species differences have been found in various mechanical properties of sclera, including compressive stiffness, tensile stiffness, and permeability [10,19,31,33,34], and this study is consistent with these findings. Here, the biggest differences between the mouse and porcine scleral properties were in the tensile stiffness, with the pig sclera being on average much stiffer than the sclera of the mouse. A minor difference in permeability was observed as well at the third strain step, with the mouse sclera being less permeable than the porcine sclera. Species differences in properties are an important factor to consider in the interpretation of results of animal models of ocular conditions. While these findings could be explained by actual species differences in the microstructure, another possibility for this study is that age differences in the two groups are driving the differences. There is ample evidence of age-related scleral remodeling of the collagen and proteoglycan networks [35–37], both of which contribute to the stiffness and permeability of the sclera. The exact age of the pigs from which the eyes were obtained was not known, so this study is limited in that this factor was not able to be considered in the model statistically as a covariate. However, the pigs were likely ∼6 months of age, compared to a mean age ∼ 4 months for the mice. This difference in mean age, especially when considering the different lifespans of the mouse and the pig, may be driving the differences in properties and how the properties change with compressive strain observed in this study. Similarly, there was a difference in strain rates applied to the sclera of the two species, which has been shown to impact measured BCLE stiffness of biphasic materials due to the intrinsic viscoelastic properties of the collagen matrix [21]. Future work will be needed to discern the exact factor driving the observed difference between the two species and whether it is an actual species difference or driven by a confounding factor. Importantly though, unconfined compression analyzed with BCLE was able to discern a difference between the two groups, showing it has the power to detect biomechanical differences between treatment groups.

### 4.7 Infinitesimal Strain Assumption

The formulation of the model used to analyze the stress relaxation experiments includes an infinitesimal strain assumption that introduces error into the analysis. While the compressive strains were held to a maximum of 15% in the stress relaxation experiments, a tare stress of ∼636Pa was applied to the samples prior to the 15% strain application. Accurately determining the corresponding amount of tare strain in these experiments is not trivial due to the curved and small size of the samples. However, the mean compressive stiffness of 10kPa for the porcine and mouse scleral samples can be used to estimate ∼6% strain due to the tare loading. While the total ∼20% compressive strain on the samples in this study introduces a nonnegligible amount of error (∼2% strain), the associated error is generally deemed to be an acceptable tradeoff for the significant simplification of the analysis in soft tissue mechanics [18,38,39]. For more objectively accurate material properties, it will be necessary to use a finite deformation formulation of the BCLE model.

## 5. Conclusion

We evaluated the applicability of unconfined compression analyzed with poroelastic theory to characterize scleral biomechanics. We showed that this methodology produces magnitudes of tensile stiffness and permeability that are consistent with those obtained by biaxial tensile testing and physiologically reasonable. Unconfined compression is moderately repeatable under a test-retest reliability experiment, and it is mostly insensitive to the thickness of the samples, as it can characterize porcine sclera as well as the significantly thinner murine sclera. Using this method, we have characterized the biomechanical properties of the murine sclera for the first time. Future work should focus on more completely validating the model’s ability to fully describe the tissue’s biomechanics by predicting another simultaneously measured quantity and work towards a full characterization of the tissue by relaxing the cubic symmetry assumption. Importantly, this method will allow increased use of the mouse as an animal model for future ocular biomechanics studies where the sclera is a target, such as myopia and glaucoma, and permit more rigorous studies of the genetic pathways regulating scleral biomechanics.

## Acknowledgements

This work was supported by the Georgia Research Alliance; NIH grants EY016435 and EY030071; Dept of Veterans Affairs Rehab R&D Service Senior Research Career Scientist Award; IK6 RX003134; and the NEI T32 Vision Research Training Grant.

## References

[1] Y. Song, F. Zhang, Y. Zhao, M. Sun, J. Tao, Y. Liang, L. Ma, Y. Yu, J. Wang, J. Hao, Enlargement of the Axial Length and Altered Ultrastructural Features of the Sclera in a Mutant Lumican Transgenic Mouse Model, (2016) 36–50. https://doi.org/10.1561/2200000016.

[2] C. Boote, I.A. Sigal, R. Grytz, Y. Hua, T.D. Nguyen, M.J.A. Girard, Scleral structure and biomechanics, Prog. Retin. Eye Res. (2019) 100773. https://doi.org/10.1016/j.preteyeres.2019.100773.

[3] R.E. Norman, J.G. Flanagan, I.A. Sigal, S.M.K. Rausch, I. Tertinegg, C.R. Ethier, Finite element modeling of the human sclera: Influence on optic nerve head biomechanics and connections with glaucoma, Exp. Eye Res. 93 (2011) 4–12. https://doi.org/10.1016/j.exer.2010.09.014.

[4] B. Coudrillier, C. Boote, H.A. Quigley, T.D. Nguyen, Scleral anisotropy and its effects on the mechanical response of the optic nerve head, Biomech. Model. Mechanobiol. 12 (2012) 941–963. https://doi.org/10.1007/s10237-012-0455-y.

[5] R. Grytz, J.T. Siegwart, Changing material properties of the tree shrew sclera during minus lens compensation and recovery, Invest. Ophthalmol. Vis. Sci. 56 (2015) 2065–2078. https://doi.org/10.1167/iovs.14-15352.

[6] J.A. Summers Rada, S. Shelton, T.T. Norton, The sclera and myopia, Exp. Eye Res. 82 (2006) 185–200. https://doi.org/10.1016/j.exer.2005.08.009.

[7] J.K. Pijanka, E.C. Kimball, M.E. Pease, A. Abass, T. Sorensen, T.D. Nguyen, H.A. Quigley, C. Boote, Changes in scleral collagen organization in murine chronic experimental glaucoma, Invest. Ophthalmol. Vis. Sci. 55 (2014) 6554–6564. https://doi.org/10.1167/iovs.14-15047.

[8] V.C. Mow, D.D. Sun, X.E. Guo, M. Likhitpanichkul, W.M. Lai, Fixed negative charges modulate mechanical behaviours and electrical signals in articular cartilage under unconfined compression a triphasic paradigm, (n.d.).

[9] Y. Yuan, M. Li, C.H. To, T.C. Lam, P. Wang, Y. Yu, X. Hu, B. Ke, The Role of the RhoA / ROCK Signaling Pathway in Mechanical Strain-Induced Scleral Myofibroblast Differentiation, (2018) 1–3.

[10] A. Eilaghi, J.G. Flanagan, I. Tertinegg, C.A. Simmons, G. Wayne Brodland, C. Ross Ethier, Biaxial mechanical testing of human sclera, J. Biomech. 43 (2010) 1696–1701. https://doi.org/10.1016/j.jbiomech.2010.02.031.

[11] B. Cruz Perez, J. Tang, H.J. Morris, J.R. Palko, X. Pan, R.T. Hart, J. Liu, Biaxial mechanical testing of posterior sclera using high-resolution ultrasound speckle tracking for strain measurements, J. Biomech. 47 (2014) 1151–1156. https://doi.org/10.1016/j.jbiomech.2013.12.009.

[12] I.C. Campbell, B. Coudrillier, C. Ross Ethier, Biomechanics of the Posterior Eye: A Critical Role in Health and Disease, J. Biomech. Eng. 136 (2014) 021005. https://doi.org/10.1115/1.4026286.

[13] K.M. Myers, F.E. Cone, H. a Quigley, S. Gelman, M.E. Pease, T.D. Nguyen, The In Vitro Inflation Response of Mouse Sclera, Exp. Eye Res. 91 (2010) 866–875. https://doi.org/10.1016/j.exer.2010.09.009.The.

[14] C. Nguyen, F.E. Cone, T.D. Nguyen, B. Coudrillier, M.E. Pease, M.R. Steinhart, E.N. Oglesby, J.L. Jefferys, H.A. Quigley, Studies of scleral biomechanical behavior related to susceptibility for retinal ganglion cell loss in experimental mouse glaucoma, Investig. Ophthalmol. Vis. Sci. 54 (2013) 1767–1780. https://doi.org/10.1167/iovs.12-10952.

[15] K. Wang, A.T. Read, T. Sulchek, C.R. Ethier, Trabecular meshwork stiffness in glaucoma, Exp. Eye Res. 158 (2017) 3–12. https://doi.org/10.1016/j.exer.2016.07.011.

[16] S. Wang, K. V. Larin, Shear wave imaging optical coherence tomography (SWI-OCT) for ocular tissue biomechanics, Opt. Lett. 39 (2014) 41. https://doi.org/10.1364/ol.39.000041.

[17] M.A. Soltz, G.A. Ateshian, A Conewise Linear Elasticity Mixture Model for the Analysis of Tension-Compression Nonlinearity in Articular Cartilage, 122 (2000).

[18] H. Hatami-Marbini, E. Etebu, An experimental and theoretical analysis of unconfined compression of corneal stroma, J. Biomech. 46 (2013) 1752–1758. https://doi.org/10.1016/j.jbiomech.2013.03.013.

[19] J.L. Battaglioli, R.D. Kamm, Measurements of the compressive properties of scleral tissue, Investig. Ophthalmol. Vis. Sci. 25 (1984) 59–65.

[20] A. Curnier, Q.-C. He, P. Zysset, Conewise linear elastic materials, J. Elast. 37 (1994) 1–38. https://doi.org/10.1007/bf00043417.

[21] C.-Y. Huang, M.A. Soltz, M. Kopacz, V.C. Mow, G.A. Ateshian, Experimental Verification of the Roles of Intrinsic Matrix Viscoelasticity and Tension-Compression Nonlinearity in the Biphasic Response of Cartilage, J. Biomech. Eng. 125 (2003) 84. https://doi.org/10.1115/1.1531656.

[22] J. Soulhat, M.D. Buschmann, A. Shirazi-Adl, A Fibril-Network-Reinforced Biphasic Model of Cartilage in Unconfined Compression, J. Biomech. Eng. 121 (1999) 340. https://doi.org/10.1115/1.2798330.

[23] Z. Hu, S. Xiong, Q. Su, X. Zhang, Sufficient conditions for global convergence of differential evolution algorithm, J. Appl. Math. 2013 (2013). https://doi.org/10.1155/2013/193196.

[24] J.R.F. Arruda, Objective functions for the nonlinear curve fit of frequency response functions, AIAA J. 30 (1992) 855–857. https://doi.org/10.2514/3.11001.

[25] W.M. Lai, V.C. Mow, Drag-induced compression of articular cartilage during a permeation experiment, Biorheology. 17 (1980) 111–123. https://doi.org/10.3233/bir-1980-171-213.

[26] T.O. Kvalseth, Cautionary Note about R2, Am. Stat. 39 (1985) 279–285.

[27] A.M. Mortazavi, B.R. Simon, W.D. Stamer, J.P. Vande Geest, Drained secant modulus for human and porcine peripapillary sclera using unconfined compression testing, Exp. Eye Res. 89 (2009) 892–897. https://doi.org/10.1016/j.exer.2009.07.011.

[28] K.S. Worthington, L.A. Wiley, A.M. Bartlett, E.M. Stone, R.F. Mullins, A.K. Salem, C.A. Guyman, B.A. Tucker, Mechanical Properties of Murine and Porcine Ocular Tissues in Compression, Exp. Eye Res. 46 (2014) 220–231. https://doi.org/10.1016/j.freeradbiomed.2008.10.025.The.

[29] F. Pennella, G. Cerino, D. Massai, D. Gallo, G. Falvo D’Urso Labate, A. Schiavi, M.A. Deriu, A. Audenino, U. Morbiducci, A survey of methods for the evaluation of tissue engineering scaffold permeability, Ann. Biomed. Eng. 41 (2013) 2027–2041. https://doi.org/10.1007/s10439-013-0815-5.

[30] T.L. Jackson, A. Hussain, A. Hodgetts, A.M.S. Morley, J. Hillenkamp, P.M. Sullivan, J. Marshall, Human scleral hydraulic conductivity: Age-related changes, topographical variation, and potential scleral outflow facility, Investig. Ophthalmol. Vis. Sci. 47 (2006) 4942–4946. https://doi.org/10.1167/iovs.06-0362.

[31] J.M. Stewart, D.S. Schultz, O. Lee, M.L. Trinidad, Permeability : Modeling the Effects of Age-Related Cross-link Accumulation, 50 (2009) 352–357. https://doi.org/10.1167/iovs.08-2300.

[32] Fluxion, Understanding effects of viscosity in the BioFlux system, Tech. Note. (2009) 1–2.

[33] B. Cruz, J. Tang, H.J. Morris, J.R. Palko, X. Pan, R.T. Hart, J. Liu, Biaxial mechanical testing of posterior sclera using high-resolution ultrasound speckle tracking for strain measurements, J. Biomech. 47 (2014) 1151–1156. https://doi.org/10.1016/j.jbiomech.2013.12.009.

[34] T.L. Jackson, A. Hussain, A. Hodgetts, A.M.S. Morley, J. Hillenkamp, P.M. Sullivan, J. Marshall, Human Scleral Hydraulic Conductivity : Age-Related Changes, Topographical Variation, and Potential Scleral Outflow Facility, (2006) 4942–4946. https://doi.org/10.1167/iovs.06-0362.

[35] J.M. Stewart, D.S. Schultz, O.T. Lee, M.L. Trinidad, Exogenous collagen cross-linking reduces scleral permeability: Modeling the effects of age-related cross-link accumulation, Investig. Ophthalmol. Vis. Sci. 50 (2009) 352–357. https://doi.org/10.1167/iovs.08-2300.

[36] J.A. Rada, V.R. Achen, S. Penugonda, R.W. Schmidt, B.A. Mount, Proteoglycan composition in the human sclera during growth and aging, Investig. Ophthalmol. Vis. Sci. 41 (2000) 1639–1648.

[37] B. Coudrillier, J. Pijanka, J. Jefferys, T. Sorensen, H.A. Quigley, C. Boote, T.D. Nguyen, Collagen structure and mechanical properties of the human sclera: analysis for the effects of age., J. Biomech. Eng. 137 (2015) 041006. https://doi.org/10.1115/1.4029430.

[38] M.A. Soltz, G.A. Ateshian, A Conewise Linear Elasticity Mixture Model for the Analysis of Tension-Compression Nonlinearity in Articular Cartilage, J. Biomech. Eng. 122 (2000) 576–86.

[39] B. Cohen, W.M. Lai, V.C. Mow, A Transversely Isotropic Biphasic Model for Unconfined Compression of Growth Plate and Chondroepiphysis, 120 (1998) 491–496.

